# Protein Kinase A Inhibition Epigenetically Silences *Ren1*

**DOI:** 10.1101/2023.09.19.558267

**Authors:** Jason P. Smith, Robert Paxton, Silvia Medrano, Nathan C. Sheffield, Maria Luisa S. Sequeira-Lopez, R. Ariel Gomez

## Abstract

**Rationale:** Renin-expressing cells are myoendocrine cells crucial for survival which detect changes in blood pressure and release renin to maintain homeostasis. One of the pathways responsible for renin expression includes cAMP as a crucial factor. cAMP binds to subunits of protein kinase A (PKA), ultimately recruiting both CBP and p300. Binding to the cAMP-responsive element in the renin enhancer region thus amplifies renin transcription.

**Objective:** To evaluate transcriptomic and epigenomic changes occurring at the renin locus via cAMP pathway inhibition.

**Methods and Results:** We treated As4.1 cells (a tumoral cell line that constitutively expresses renin) with the PKA inhibitor H89 (treated) or DMSO (control). We then performed independent ATAC-seq, scRNA-seq, and ChIP-seq for H3K27Ac and P300 binding on biological replicates of treated and control As4.1 cells. *Ren1* expression is significantly reduced following PKA inhibition with a corresponding loss in H3K27Ac and P300 binding at the locus. A restricted set of nine genes with overlapping dynamically accessible regions, differential gene expression, and H3K27Ac and P300 binding were identified with roles among three primary renin regulatory paradigms.

**Conclusions:** The data suggests that cAMP pathway inhibition controls renin expression through a reduction not in accessibility alone, but via a switch from an active to poised state of epigenetic control, a shift towards a less differentiated cellular identity, and the disruption of not only cAMP, but baroreceptor and Notch mediated renin regulatory pathways.

## Introduction

Renin is a hormone enzyme that is the rate-limiting component of the Renin-Angiotensin Aldosterone System (RAAS or RAS). It is critical to our survival through the maintenance of homeostasis and is predominantly produced in a rare subset of cells located in the kidney next to glomeruli known as juxtaglomerular (JG) cells. Upon threats to homeostasis, descendants of the JG cell progenitors, including vascular smooth muscle cells, mesangial cells, and interstitial pericytes, may revert to a renin-expressing phenotype. As this phenotype switching role is likely regulated at both the genetic and epigenetic level, how exactly cells turn off or on the renin phenotype is of upmost importance to our understanding of the control of homeostasis.

While the rarity of JG cells (Castellanos-Rivera et al., 2015) and their intractability to culture (Karginova et al., 1997) stymie many classical experimental methods, much of the pathways and molecular signatures that regulate renin production have been revealed. Renin regulation is modulated by three primary signaling pathways: cAMP signaling, Notch signaling, and through a baroreceptor mechanism. Here, we focused on the cAMP pathway where the binding of the cAMP-responsive element (CRE) in the renin enhancer region amplifies renin transcription and is the most important driver of renin synthesis (Castrop et al., 2010). This occurs through a mechanism where cAMP binds to subunits of protein kinase A (PKA) that ultimately recruits both CBP (CREB-binding protein) and the histone acetyl-transferase p300 to the renin locus. Because cells at the juxtaglomerular apparatus and along the arterioles can transiently alter their renin phenotype, their chromatin must be poised to enable such changes to occur and supports a conservation of transcriptional machinery present in all JG progenitor cell descendants.

While it is known that disruption of PKA will reduce *Ren1* abundance (Pan et al., 2001), the specific gene signatures and chromatin changes that occur following such treatment remains unclear. Thus, the aim of the current study was to improve our understanding of the complex interactivity among renin regulatory pathways. To do so, we disrupted the upstream effector of the cAMP pathway, PKA, and evaluated the genetic and epigenetic shifts that occurred using a combination of ATAC-seq, scRNA-seq, and ChIP-seq experiments. Due to the inherent challenges in studying JG cells as outlined above, we treated As4.1 cells (a tumoral cell line that constitutively expresses renin) with the potent PKA inhibitor H-89-dihydrochloride (H89) or DMSO as a control and evaluated the resultant genetic and epigenetic changes.

## Methods

### Cell culture

The renin-expressing As4.1 cell line (ATCC, CRL-2193) (Sigmund et al., 1990) was maintained in high glucose DMEM (supplemented with 10% fetal bovine serum (both from Thermo Fisher Scientific) at 37°C in a humidified incubator containing 5% *CO*_2_. Biological replicates (n=2) for each experiment (ATAC-seq, scRNA-seq, and H3K27Ac and P300 ChIP-seq) were independently passaged and treated with either a control (DMSO) or the PKA inhibitor, H-89-dihydrochloride (H89, Cell Signaling 9844S), at 0-, 24-, and 48-hours following experiment initiation. After 48 hours, cells were subjected to bulk ATAC-seq, scRNA-seq, or ChIP-seq for both H3K27Ac and P300.

### ATAC-seq

ATAC-seq (Buenrostro et al., 2013) was performed on cultured As4.1 cells treated with either H89 (n = 2) or DMSO (n = 2) at 0, 24, and 48 hours. We collected 50,000 cells, washed them 3X with 5 mL dPBS (Dulbecco’s Phosphate Buffered Saline, Gibco, UK), treated them with Lysis Buffer, and resuspended them and performed tagmentation using a Nextera DNA library Prep Kit (Illumina). Paired-end ATAC-seq libraries were sequenced by Novogene (Durham, NC).

### ATAC-seq Read Preprocessing

For intial processing of sequenced reads, the 150 bp paired-end reads were aligned to the mouse genome (GRCm38/mm10) using the PEPATAC (v0.10.4) pipeline (Smith et al., 2021). Enzymatic bias of aligned reads was corrected using seqOutBias (Martins et al., 2017) prior to the merging of biological replicates. Peak calling on merged replicates was performed using Model-based Analysis for ChIP-Seq 3 (Zhang et al., 2008) with the following parameters `-B --call-summits --keep-dup 50 -q 0.05 -m 10 200`. The summits of peaks were extended 100 bp up and dowstream of the summit, blacklisted sites removed, and only those peaks with a -log10(pvalue) greater than 6 were retained for downstream analysis. All samples were also merged, and a consensus peak set generated using the same parameters as above.

### ATAC-seq Differential Peak Analysis

Differentially accessible regions were identified using DESeq2 (Love et al., 2014), and we clustered those dynamic peaks using DEGreport (Lorena Pantano, 2023). De novo motif identification on the clusters of dynamic peaks was evaluated using MEME (Bailey et al., 2015a) and TOMTOM (Gupta et al., 2007) against a combined motif database from HOMER (Heinz et al., 2010), JASPAR (Castro-Mondragon et al., 2021), uniPROBE (Hume et al., 2015), and cis-bp TF databases (Weirauch et al., 2014). FIMO (Grant et al., 2011) was then employed to identify the composite motif occurrences present in peaks genome wide. Finally, the bigWig (https://github.com/andrelmartins/bigwig, v0.2.9) package was used to evaluate motif enrichment around the summits of ATAC-seq regions. Dynamic region enrichment analysis was performed on the individual dynamic peak classes using GREAT Tanigawa et al. (2022).

### scRNA-seq

We plated 600,000 cultured As4.1 cells in T25 flasks, in duplicate. The cells were treated with 10 *μ*M H89 or equivalent volume DMSO at 0, 24, and 48 hours. After 48 hours, cells were trypsinized and 4000 cells per condition were targeted for sequencing. The cells were centrifuged in a Sorvall RT7 refrigerated 4°C centrifuge (Sorvall, Newtown, CT) at 500 x g for 10 minutes, washed 3X with 5mL of dPBS, then 1 mL of trypsin was added and cells incubated at 37°C for 2 minutes before adding 4 mL of fresh media. The cells were gently pipetted up and down 20 times to dissociate and centrifuged at 800 x g for 3 minutes at room temperature. The supernatant was removed, and pellets resuspended in 1.5mL dPBS supplemented with 0.04% BSA (Ultra-PureTM Bovine Serum Albumin, Invitrogen, Lithuania) by pipetting up and down 20 times. Resuspended cells were passed through a 40 *μ*m filter and repelleted by centrifugation for 3 minutes at room temperature at 150 x g. The supernatant was carefully removed, and cells washed 3X in 1.5mL dPBS with 0.04% BSA as outlined above. Cells were resuspended a final time in 1.5mL dPBS with 0.04% BSA and the volume adjusted to yield 1000 cells/*μ*L. Single cells were then loaded and captured using the Chromium System (10X Genomics, Pleasanton, CA) following the manufacturer’s recommendation (Magaletta et al., 2022) for the Chromium Next GEM Chip G with reagents of Chromium Next GEM Single Cell 3^′^ Reagent Kits v3.1 (10X Genomics). Finally, the completed libraries were sent for sequencing by Novogene (Durham, NC).

### Read preprocessing

#### Genome and transcriptome annotations

We performed scRNA-seq analyses using the mm10 reference. For the initial read processing, we employed the Cell Ranger `refdata-gex-mm`10-2020-A reference data and for analysis in the R programming language, the `BSgenome package` `BSgenome.Mmusculus.UCSC.mm10`.

#### scRNA-seq alignment and feature-barcode matrix generation

scRNA-seq fastq files from the 10X Genomics Single Cell Gene Expression platform were processed using the Cell Ranger pipeline (version: `cellranger-7.1.0`). This pipeline trims non-template sequence and aligns reads with the STAR aligner (Dobin et al., 2013). Following mapping, transcriptomic reads are grouped by barcode, UMI, and gene with reads mapped to genes calculated using UMI-based counts. The filtered UMIs were linked to barcodes and the resultant feature-barcode matrices were employed for downstream analysis.

#### scRNA-seq quality control

Feature-barcode matrices produced by Cell Ranger were analyzed in R using Seurat (Stuart et al., 2019). Cells with mitochondrial (mt-) or hemoglobin (Hb-) DNA less than the 97.5^th percentile were retained for further analysis. Retained cells were required to include the number of identified RNA features between the 2.5^th and 97.5^th percentiles, as extremely low or high numbers of features are indicative of low quality or doublet cells (McCarthy et al., 2017; Ziegenhain et al., 2017).

#### scRNA-seq cell clustering

Cells were normalized and variance-stabilized using a regularized negative binomial regression implemented in R (see Seurat SCTransform), while controlling for the cell cycle, mt-DNA, and Hb-DNA percentage (Hafemeister and Satija, 2019; Stuart et al., 2019). We integrated the DMSO and H89 treated cells by selecting the top 3000 most variable features (see `Seurat SelectIntegrationFeatures and IntegrateData`). Principal Component Analysis (PCA) and Uniform Manifold Approximation and Projection (UMAP) were run on the normalized data prior to the contruction of a shared nearest neighbors graph to identify cell clusters (see Seurat `RunPCA, RunUMAP, FindNeighbors, and FindClusters`).

#### scRNA-seq cell identification

The top 25 differentially expressed genes present in at least 25% of the cells in a cluster and with a minimum 0.25 log fold change between clusters (see Seurat `FindAllMarkers`) were defined as markers for each cluster. These genes were used to annotate individual clusters. Because the single-cells here are derived from a single cell-line, clusters were annotated by the presence, or absence, of specific markers over- or underrepresented in each cluster.

### Gene set enrichment analysis (GSEA)

We evaluated the enrichment of annotated gene sets from the Molecular Signatures Database for mouse hallmark gene sets, regulatory target gene sets, curated gene sets, and cell signature gene sets. This procedure was performed in `R` using the `fgsea` package for the following comparisons: all DMSO (control) cells and all H89 (treated) cells, and for the same comparison within each identified cell cluster.

### Velocity analysis

We performed velocity analysis of the scRNA-seq interrogated cells to study the cellular dynamics following PKA inhibition via H89 treatment. The Cell Ranger output directory was used as input to the `Python` package `velocyto` (La Manno et al., 2018) to generate spliced and unspliced expression matrices. The spliced and unspliced matrices were then added as additional features of the `Seurat` objects in R and the `RunVelocity` function from the `SeuratWrappers` package was used to visualize the RNA dynamics on the `Seurat` embeddings.

### ChIP-seq

We plated 4.2 million As4.1 cells (passage 29) per 150 mm plate, in duplicate. Cells were treated with 10 *μ*M H89 or equivalent volume DMSO at 0, 24, and 48 hours. Cells were then fixed according to manufacturer protocol prior to shipping to Active Motif (Carlsbad, CA) for sequencing. Chromatin was precipitated with antibodies against H3K27ac and P300. ChIP and Input DNAs were sequencing using a NextSeq 500 (Illumina) and the 75 nt raw reads were processed using the `nf-core/chipseq` pipeline (Ewels et al., 2020) with alignment to the GRCm38/mm10 mouse genome. Enriched genomic regions for both H3K27Ac and P300 were determined using MACS2 (Model-based analysis of ChIP-seq 2) (Zhang et al., 2008) with default settings by comparing ChIP to input samples. Differential binding analysis was then performed using the `DiffBind` package `Ross-Innes` et al. (2012). Differentially enriched regions were annotated using the `ChIPpeakAnno` Zhu (2013) and `clusterProfiler` Wu et al. (2021) packages.

### Integrative Analysis

To identify the specific subset of genomic regions and associated target genes disrupted by PKA inhibition, we first overlapped the two dynamic peak clusters (either increasing or decreasing accessibility) with the differentially expressed genes identified in the `“Ren1 LO”` subcluster of the scRNA-seq experiment. With two subsets of regions and target genes identified, we next identified which of those region-gene sets also overlapped differential H3K27Ac and P300 binding sites. The resultant subsets revealed a minimal set of genes with overlaps to either the dynamically increased accessibility regions or dynamically decreased regions.

## Results

### ATAC-seq Reveals Distinct Clusters of Dynamically Accessible Chromatin Following PKA Inhibition

We identified more than 120000 accessibility peaks with a small fraction representing dynamically accessible regions (n = 12260) following PKA inhibition. We clustered those dynamic peaks on their kinetic profiles and identified two region classes: one where the clustered regions increased (n = 1959) in accessibility upon PKA inhibition, and a second cluster which decreased (n = 1106) in accessibility following treatment (Fig. 1A). Within the decreased accessibility region cluster, these putative regulatory elements (REs) were restricted primarily to intronic and distal intergenic regions, implying a greater abundance of enhancer-like elements losing accessibility following treatment (Fig. 1B, top panel). Conversely, the increased accessibility cluster were found to contain promoter-centric elements (30.4%, 1B, lower panel) located primarily in 5’ UTRs (Fig. 1B). Therefore, PKA inhibition displays disparate effects on the category of disrupted RE, with promoter-like REs more likely to increase in accessibility and enhancer-like REs losing access. Enhancers are associated with specific cell identity and function (Heinz et al., 2015), and here we observe a loss in accessibility specifically within the superenhancer integral to *Ren1* expression (Fig. 1C).

**Fig. 1:**
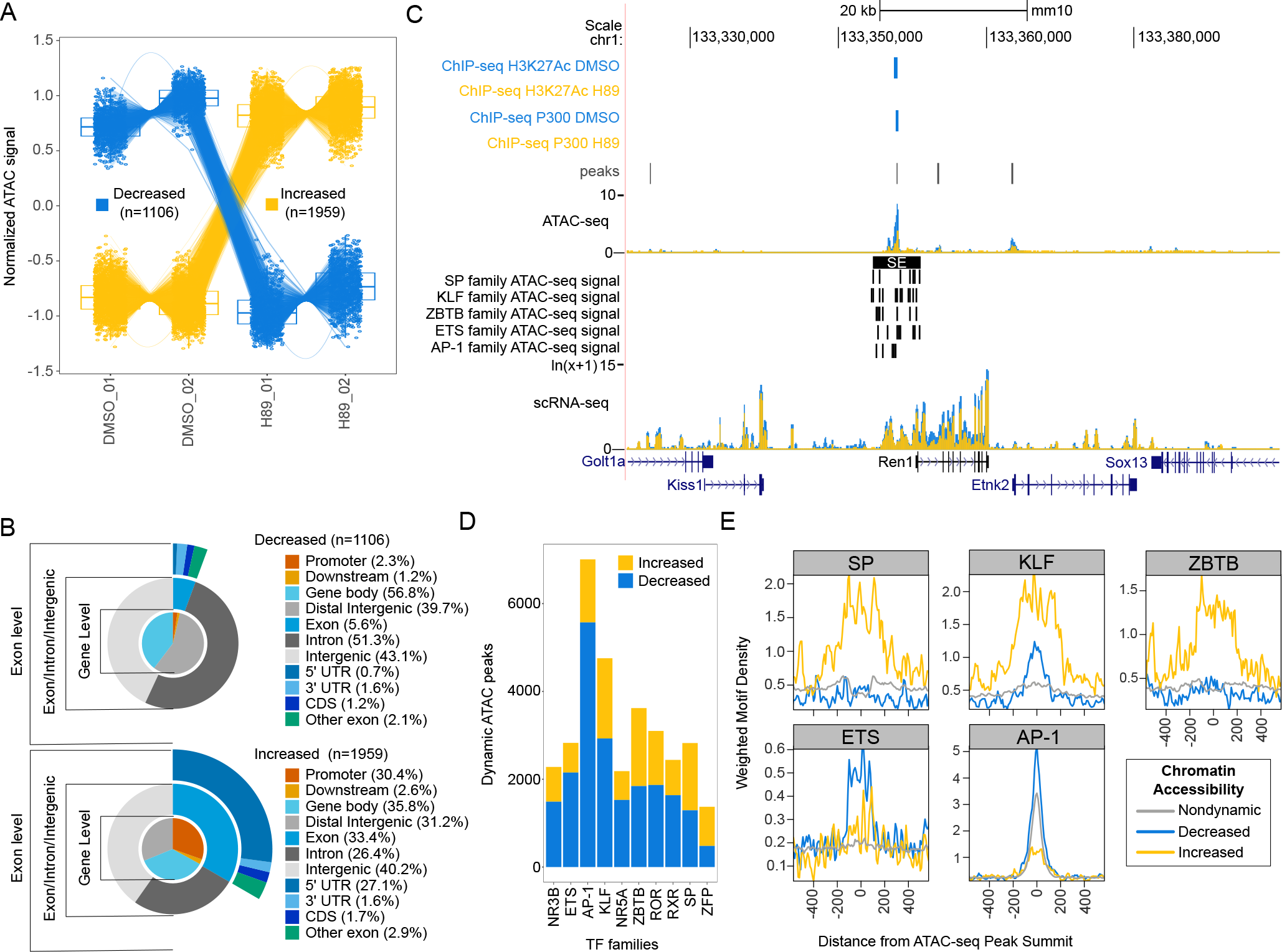
Chromatin analysis identifies loss of Ren1 accessibility and SP/KLF and ZBTB transcription factors as important regulatory features. (A) Dynamic peak clusters based on kinetic profiles. (B) Genomic distribution of dynamic peak clusters. (C) Ren1 locus with binding profiles for H3K27Ac, P300, open chromatin, gene expression, and identified transcription factor (TF) families. (D) The distribution of dynamic peak clusters among enriched TF families. (E) Composite motif densities for the TF families overrepresented in the two dynamic peak clusters.

Next, we performed de novo motif analysis (Bailey et al., 2015b) within the dynamic peaks and identified 10 primary TF family motifs (Fig. 1D). Of these TF families, the SP, KLF, and ZBTB families are overrepresented in the increased accessibility dynamic peak cluster (Fig. 1E). SP and KLF members are reported to serve a developmental role in the basal transcription of renin and bind to the proximal promoter region of the *Ren1* locus (Todorov, 2013). SP family members, specifically *Sp1*/*Sp3*, bind at *Ren1* and promote transcription (Pan et al., 2003), although the locus itself is less accessible following PKA inhibition (Fig. 1C), suggesting the action of SP members occurs at alternate target genes. In support, these two SP members, *Sp1* and *Sp3*, are the highest expressed members of the family (Supplemental Fig. S1A). KLF members, with reported binding sites at *Ren1* (Martinez et al., 2018), associate with several renal physiological characteristics including: endothelial cell barrier function and damage prevention, antiapoptosis in podocytes, and renal fibrosis (Rane et al., 2019). Within this dataset, *Klf4, Klf3*, and *Klf13* are the highest expressed family members (Supplemental Fig. S1A).

ZBTB members are important regulators of chromatin interactions in conjunction with cohesin (Wang et al., 2023), emphasizing that physical changes in chromatin structure occur following PKA inhibition that abrogate the ability of the cells to continue expressing renin. *Zbtb20* is the highest expressed ZBTB member (Supplemental Fig. S1A), with previous studies pointing to *Zbtb20* dysregulation playing a role in hypertensive renal tissue although the exact mechanisms remain unclear (Wei et al., 2023).

The ETS and AP-1 families are overrepresented in the decreased accessibility peak cluster (Fig. 1E). ETS members, in particular *Ets1*, can promote renin expression through binding at the gene locus Martini et al. (2023). Here, ETS members *Elk3, Etv6*, and *Elf1, Elf2*, and *Elf4*, are the highest expressed members (Supplemental Fig. S1B). AP-1 has long been known to regulate renin expression (Tarnura et al., 1992), and interactions between ETS and AP-1 have recently been associated with an aging phenotype in the kidney (Yu et al., 2023). In the companion scRNA-seq data, *Jund, Maf*, and *Junb* are the highest expressed AP-1 members (Supplemental Fig. S1B) with *Jund* known to be a part of the cAMP pathway (Martinez et al., 2018) and involved in *Ren1* regulation (Martini et al., 2023).

### scRNA-seq Reveals Subset of Cells with Significantly Altered Ren1 Expression Following PKA Inhibition

To evaluate the effect of PKA inhibition, we also performed scRNA-seq on DMSO or H89 treated As4.1 cells. As expected and previously reported (Pan et al., 2001), *Ren1* expression is significantly ({T_Welch’s_} *ρ*=0.00) reduced with PKA inhibition (Fig. 2A). We integrated the scRNA-seq of the H89 treated (trt) and DMSO treated (ctrl) cells (Fig. 2B), and identified six cell clusters with distinct gene signatures (Fig. 2C). The largest cluster expressed a combination of every marker composing the additional clusters, and therefore we termed it “all+”. The “remodel” cluster included several histone family members (e.g., *Hist1h2ap, Hist1h1b, Hist1h1e*) as the most significant markers, with an overall enrichment of cell proliferative pathways (Supplemental Fig. S2). The “*Khdrbs3*|*Acaa1b*+”, “*Sec24a*|*Arid3b*+”, and “*Gbp7*+” clusters overexpressed their named markers relative to the individual cell populations. Finally, the “*Ren1* LO” cluster represented the population of single cells with the lowest expression of renin (Supplemental Fig. S2 A). Next, we generated spliced and unspliced transcript matrices to investigate RNA dynamics (La Manno et al., 2018).

**Fig. 2:**
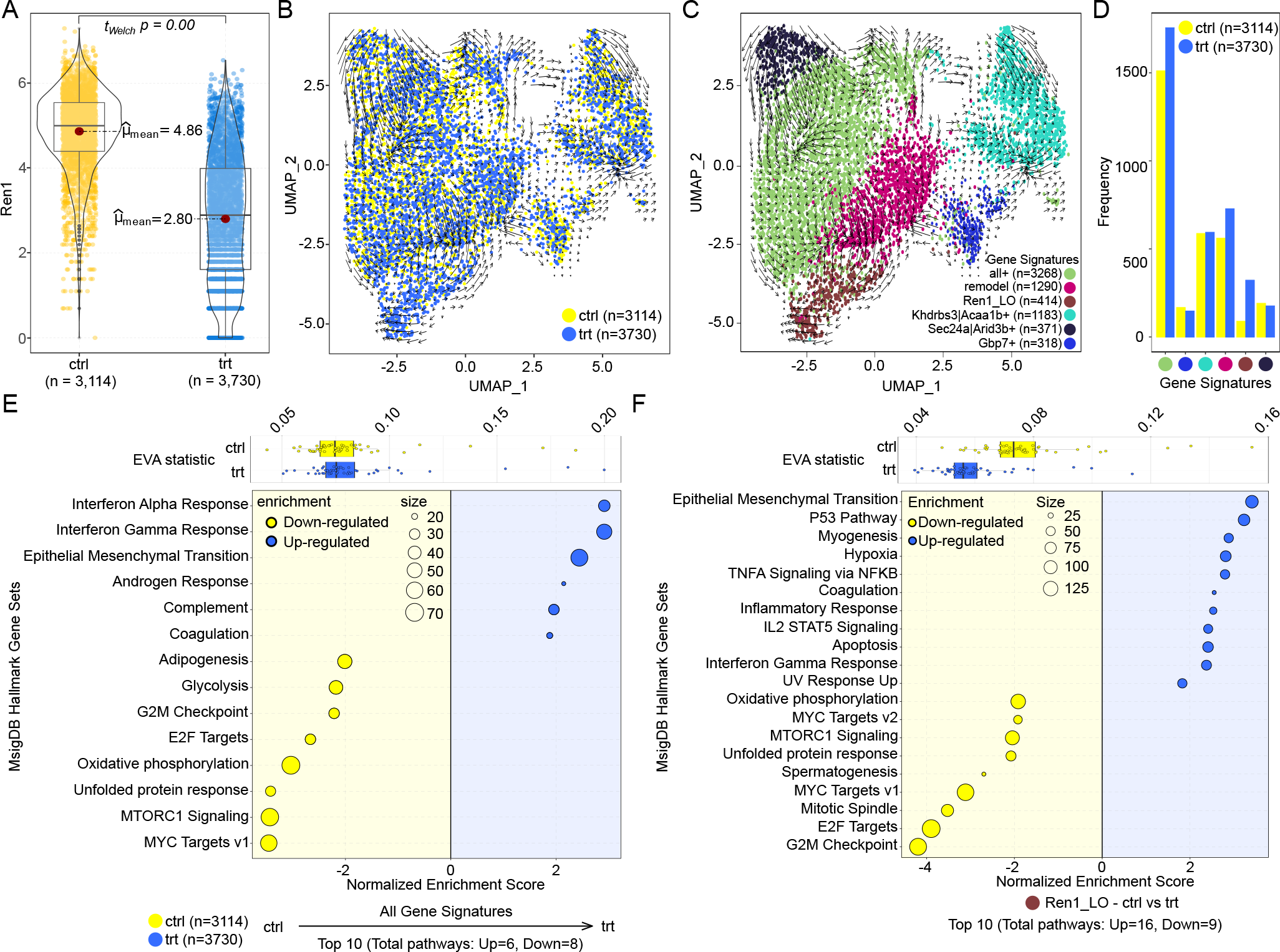
scRNA-seq reveals changes in RNA dynamics and pathway heterogeneity. (A) Ren1 expression in PKA inhibited (trt) or DMSO-treated (ctrl) cells. (B) scRNA-seq UMAP with RNA dynamics colored by experimental condition. (C) scRNA-seq UMAP with RNA dynamics colored by annotated cell population gene signatures. (D) The number of single cells of each experimental condition separated by population gene signatures. (E) The hallmark gene sets and EVA statistics for the entire data set comparing trt and ctrl. (F). The hallmark gene sets and EVA statistics for the “Ren1 LO” cluster comparing trt and ctrl.

As the reduction in *Ren1* expression was our target via PKA inhibition, we investigated the “Ren1 LO” cluster in greater detail to identify which genes and pathways were disrupted. We observed the RNA dynamic vector field and identify movement away from the “Ren1 LO” cluster, suggesting lowered *Ren1* expression is a transient cell state leading to alternative, stable phenotypes of altered cell identify and function in the other clusters (Fig. 2B, C). Further, this cluster contained the highest trt to ctrl ratio of cells (Fig. 2D), suggesting that *Ren1* expression is preferentially depleted, as expected, in PKA inhibited cells. To evaluate which pathways are enriched, we evaluated the relative heterogeneity (Davis-Marcisak et al., 2019) and performed gene set enrichment analysis (GSEA) (Korotkevich et al., 2021) between trt and ctrl cells at the data set level and within each cluster (Fig. 2D, E and Supplemental Fig. S2B). In this context, heterogeneity is defined as differentially variable pathways between the PKA inhibited or DMSO treated cells. At the full data set level, there is no difference in pathway heterogeneity between trt and ctrl (Fig. 2E, F). However, within the “Ren1 LO” cluster of interest there is a noticeable reduction in heterogeneity among the PKA inhibited cells. This suggests PKA inhibition directs cells toward a more homogenous state with decreased transcriptional complexity, tying into the reduction in distal RE accessibility and a promoter-centric, less differentiated cell identity as seen in the ATAC-seq data. This is further reflected via enrichment analysis, with proliferative programs (including: E2F targets, GSM checkpoint, and MYC targets) being repressed in the PKA inhibited cell populations and an epithelial to mesenchymal transition promoted (Fig. 2E, F, Supplemental Fig. S2B).

### ChIP-seq Identifies Specific Reduction in H3K27Ac and P300 Binding at the *Ren1* Locus

We next performed ChIP-seq for both H3K27Ac and p300 binding in DMSO or H89 treated As4.1 cells. At the *Ren1* locus specifically, we observe the differential loss of H3K27Ac and p300 binding within the *Ren1* superenhancer (Fig. 1B). Together with the observation of a reduction in accessibility and expression at the locus, we find a repressed gene locus with a switch from an active state of transcription to a likely poised state of activity (Creyghton et al., 2010), although we would need to include H3K4me1 and H3K27me3 in the future to confirm a poised versus fully repressed state. However, as p300 binding represents a primary activator of *Ren1* transcription and we observe its loss at *Ren1* in combination with a reduction in accessibility through changes in chromatin structure supported by the activity of ZBTB TFs, the data points toward a switch from active to poised transcription where an observed, reduced expression of *Ren1* still occurs opposed to full repression (Fig. 2A).

Broadly, PKA inhibition led to a reduction in distal H3K27Ac and an increase in the distribution in promoter regions (Supplemental Fig. S3A, top). As H3K27Ac at enhancer regions is correlated to target gene expression (Creyghton et al., 2010), the shift from distal to proximal regions identifies a global loss in cell identity dependent gene activation in PKA inhibited As4.1 cells. Conversely, the P300 binding profile is the inverse (Supplemental Fig. S3A, bottom), with PKA inhibition leading to a broader regional distribution. As the acetyl-transferase p300 is associated with enhancers (Ghisletti et al., 2010), and enhancers are associated with cell identity, the shift in p300 binding represents a cell-wide programmatic shift in cellular identity stemming from PKA inhibition. Further, the ETS protein PU.1 has been reported to be required for maintaining H3K4me1 at p300 bound enhancers (Ghisletti et al., 2010), and we identified overrepresented ETS motifs within the dynamic regions with decreased accessibility following PKA inhibition (Fig. 1E). This supports a switch from active to poised states of transcription in genes required for *Ren1* expression in untreated As4.1 cells.

### Integrated Analysis Identifies a Subset of Genes Related to All Known Renin Regulation Pathways

Next, we integrated findings across each of the above -omics data to identify the specific subset of chromatin regions and genes most affected by PKA inhibition. Across the chromatin regions with increased accessibility, we identified those regions’ nearest differentially expressed genes with overlapping differential H3K27Ac and P300 sites following PKA inhibition. This revealed a restricted set of gene targets most affected by PKA inhibition which included the following genes: *Asap1, Tbc1d2, Rsu1, Tm4sf1, Btg1, Ndrg4, Anxa4, Il17ra*, and *Scd1* (Supplemental Fig. S4).

*Asap1* plays a role in mesenchymal progenitor cell differentiation (Schreiber, 2019), regulates integrin beta-1 recycling (Onodera et al., 2012), and plays a role in chromatin remodeling related to cellular growth regulation (Brown et al., 1998). Intriguingly, recent work identified the renal baroreceptor mechanism as residing within renin-expressing cells, with signal transduction requiring integrin beta-1 at the cell membrane (Watanabe et al., 2021). Coinciding with the role of *Asap1* in integrin beta-1 recycling, *Tbc1d2* is also identified within this subset and participates in cellular trafficking, specifically by modulating *Rab7* positive endosomes (Raudenska et al., 2021) which recycle integrin beta-1 (Margiotta et al., 2017). The involvement of integrin recycling and activity is further emphasized by the involvement of *Rsu1* and *Tm4sf1. Rsu1* interacts indirectly with integrin beta-1 via the PINCH/ILK complex, ultimately inhibiting NF-κB. *Tm4sf1* interacts constitutively with integrin beta-1 (Shih et al., 2009) and can promote AKT phosphorylation (Rahim et al., 2023).

Both *Btg1* and *Ndrg4* also interact with *Akt* signaling. *Btg1* is a mediator of cellular stress signaling pathways (Yuniati et al., 2019) with the ability to inhibit the PI3K/AKT signaling pathway (Zhang et al., 2019). *Ndrg4* is known to promote the transcription of myogenic factors via AKT and CREB activation with the reduction of *Ndrg4* leading to a decrease in CREB binding (Zhu et al., 2017). Further, the myogenic factor myocardin (*Myocd*) is active in renin-expressing cells through the promotion of smooth muscle cell genes (Brunskill et al., 2011), which are themslves descendants of renin progenitors in the kidney and are known to rexpress renin under homeostatic threat. Here, the concomitant increase in *Ndrg4* expression would promote CREB activity, but with the corresponding loss in accessibility at *Ren1*, alternate AKT/CREB targets would correspondingly be favored. Indeed, we see significant EMT pathway enrichment in PKA inhibited cells (Fig. 2E, F), which is a known outcome of disrupted CREB signaling (Mehra et al., 2022). *Anxa4* is also linked to increased AKT phosphorylation and is mediated by intracellular Ca^2+^ levels (Lin et al., 2012). As we are specifically inhibiting PKA, and PKA can inhibit calcium channel inhibitors themselves (Papa et al., 2022), the reduction in PKA activity should increase intracellular calcium, thereby stimulating *Anxa4*. Additionally, increased intracellular calcium would expectingly lead to further *Ren1* reduction, as increased intracellular Ca^2+^ is known to inhibit renin synthesis (Atchison and Beierwaltes, 2013). Providing yet more negative feedback on cAMP driven stimulation of *Ren1, Anxa4* is a direct inhibitor of adenylyl cyclase type 5 Heinick et al. (2015), thereby acting upstream of PKA inhibition. Finally, *Anxa4* can also suppress NF-κB transcriptional activity (Jeon et al., 2010).

The link to NF-κB can be further supported through the activity of both *Il17ra* and *Scd1. Il17ra* can activate both NF-κB and MAPK pathways (Hata et al., 2002), with MAPK being necessary for PKA coupling to CREB phosphorylation under normal cellular conditions (Roberson et al., 1999). *Scd1* interacts with NF-κB in a positive feedback loop (Li et al., 2017), and the overactivation of NF-κB signaling can promote stemness-associated and inflammatory gene programs (DiDonato et al., 2012).

## Discussion

Regulation of renin is well established to be modulated by many complicated mechanisms involving genetic and epigenetic control and the interaction of the cAMP, Notch pathways, and through mechanical transduction of pressure-mediated changes via a baroreceptor (Fig. 3A).

**Fig. 3:**
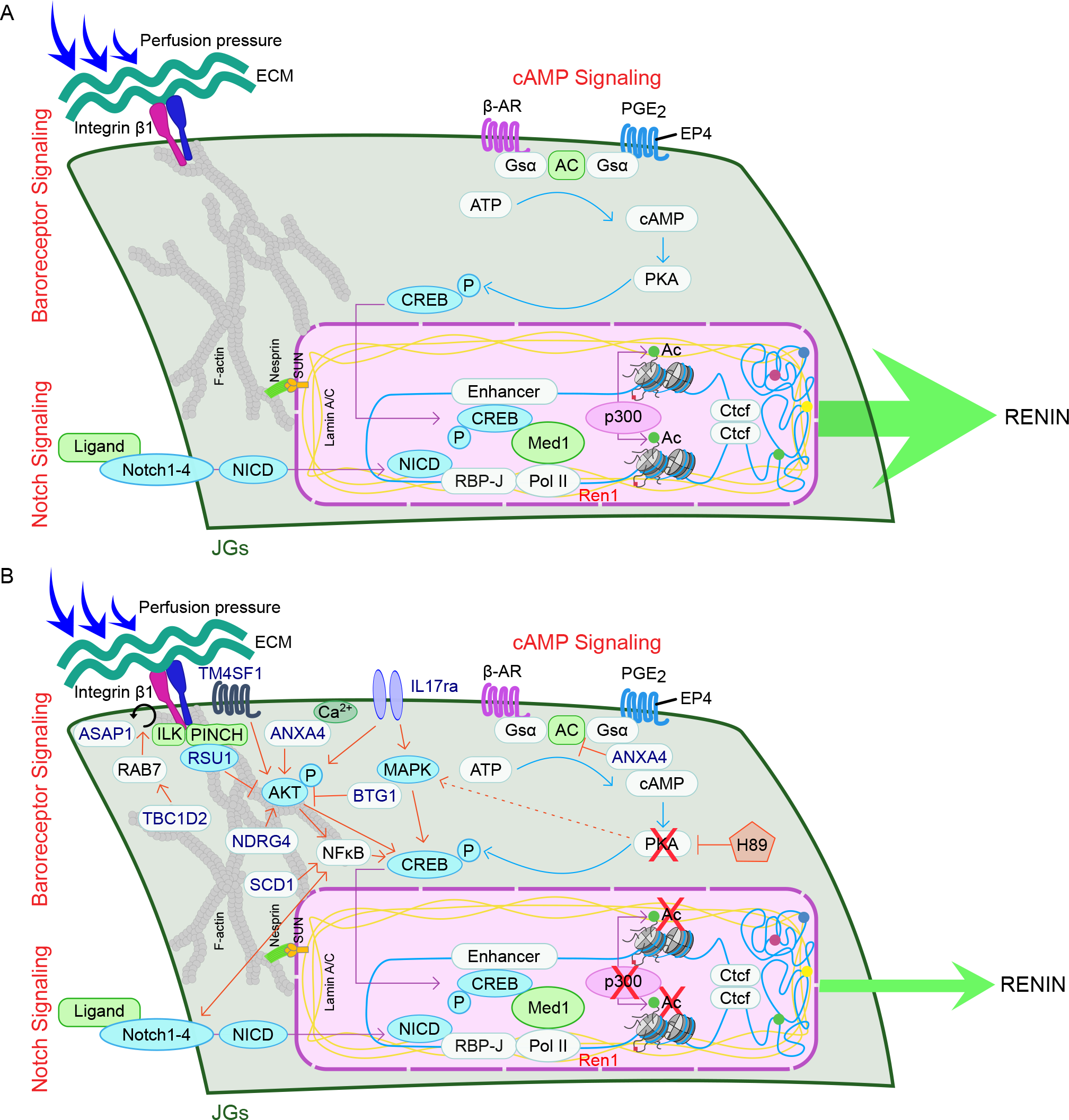
PKA inhibition represses Ren1 transcription and disrupts cAMP, baroreceptor, and Notch mediated signaling pathways. (A) The steady-state Ren1 locus. (B) The PKA inhibited Ren1 locus.

The restricted set of genes with increased expression, accessibility, H3K27Ac, and P300 binding coincide with an epithelial-to-mesenchymal transition (EMT) and a more stem-like phenotype. This phenotype would coincide with the combinatorial activation of AKT (Yoon et al., 2021), NF-κB (Taniguchi and Karin, 2018), and MAPK signaling (Huang et al., 2016) (Supplemental Fig. S5). AKT is critical for the maintenance of mesenchymal stem cells while NF-κB promotes genetic and epigenetic alterations that promote EMT and stem cell properties, in addition to interaction with a number of additional signalling pathways, including Notch (Ferrandino et al., 2018).

Both NF-κB and PPARγ are also directly or indirectly involved in the cAMP pathway in renin-expressing cells. There are known interactions between NF-κB and Notch (another major driver of renin expression) that leads to increased expression of Notch target genes, including *Ren1*, but also *Hes1*, which suppresses PPARγ expression (Maniati et al., 2011). PPARγ can directly bind to *Ren1* to promote transcription and it indirectly stimulates the cAMP pathway (Todorov, 2013), which in this context is already abrogated through inhibition of PKA.

Therefore, a loss of PPARγ would be expected to lead to a reduction in renin activation along both direct and indirect pathways. Further, NF-κB can interact with the renin locus either through direct binding at a consensus sequence in the renin enhancer, or through a competitive binding of the p65 subunit of NF-κB at the renin cAMP response element (CRE) (Todorov et al., 2005). Therefore, attenuated renin transcription can occur through the competitive blocking of CREB binding at the renin CRE. Therefore CREB, along with PPARγ (Todorov, 2013) and NF-κB, regulate renin in response to homeostatic threat or other physiological signals through an interaction within the enhancer of *Ren1*.

PKA inhibition leads to complex interactions among all of the known renin regulating pathways pushing the cell towards a more stem-like, motile phenotype. Although we are specifically disrupting the cAMP pathway, the most affected genes are tied to reductions not only in cAMP-related pathways, but in cell surface integrin expression affecting the baroreceptor mechanism, intracellular Ca^2+^ mediated regulation, and Notch signaling via NF-κB (Fig. 3B).

## Perspectives and Significance

Here, we presented a combination of three techniques, ATAC-seq, scRNA-seq, and ChIP-seq that combined reveal a more complex and nuanced understanding of the gene signatures and chromatin disruptions that occur following PKA inhibition in renin expressing cells. While the specific direct roles of the identified proteins in renin regulation remain to be further elucidated, this work provides a first look at the specific gene pathways that are altered with tie ins to all the known pathways of renin regulation. This emphasizes the complex interplay of regulation at the genetic and epigenetic level and future efforts could target these reported genes to better understand how a cell may turn off or on the renin expressing phenotype.

## Supplemental figures

**Fig. S1:**
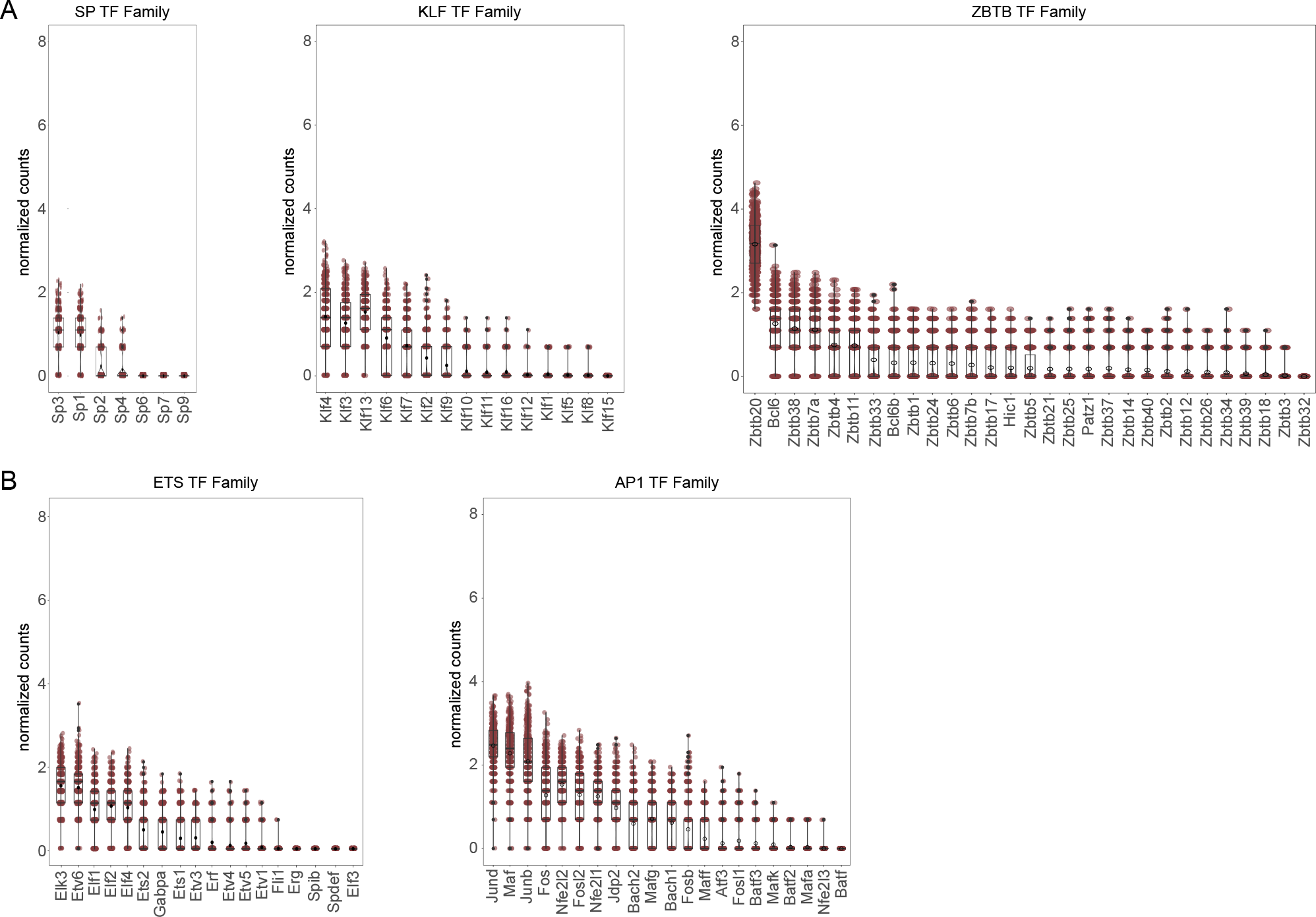
Normalized expression of overrepresented transcription factor (TF) family members. (A) Gene expression for TF family members overrepresented in regions with **increased** accessibility. (B) Gene expression for TF family members overrepresented in regions with **decreased** accessibility.

**Fig. S2:**
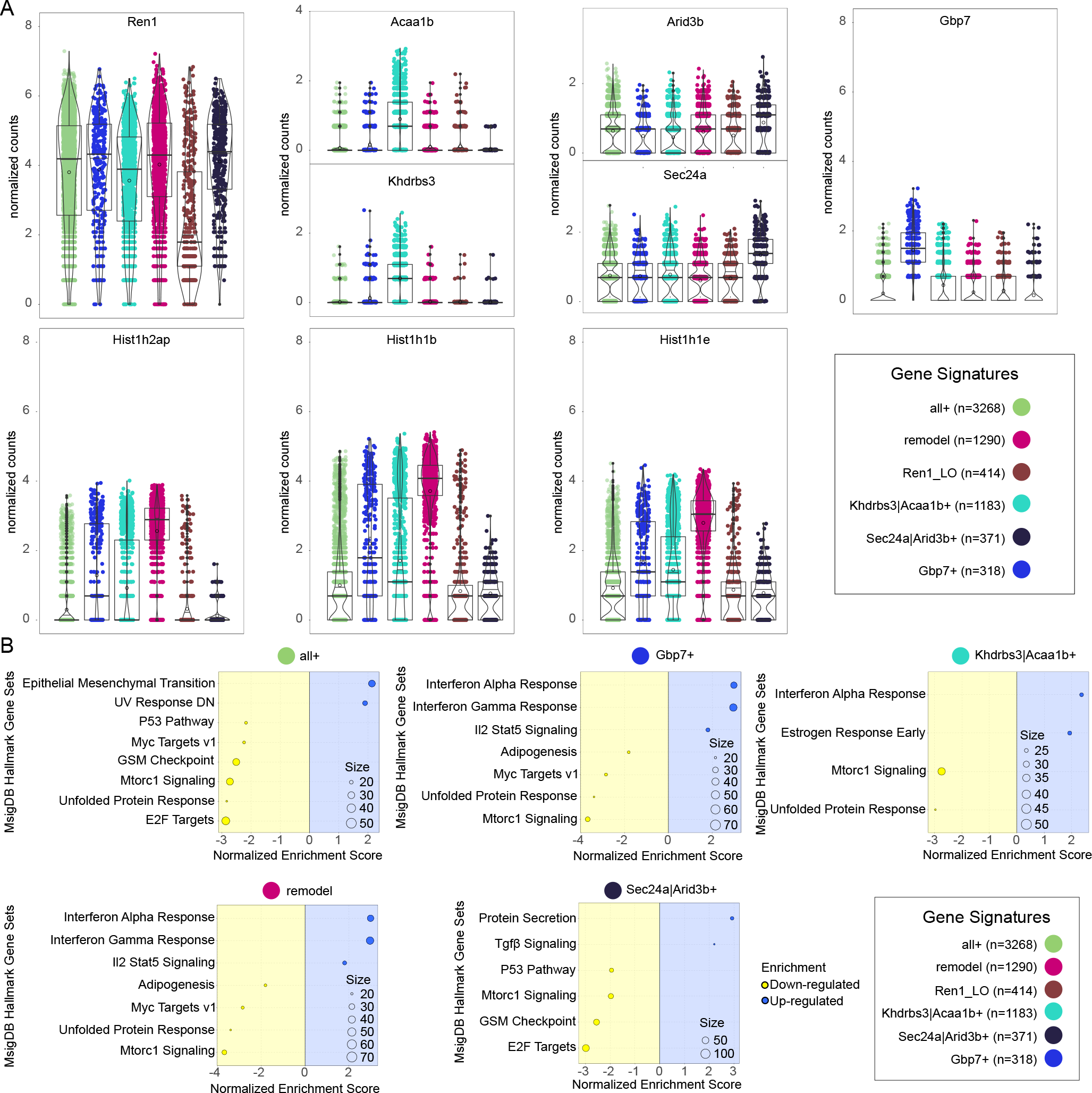
Distinct scRNA-seq clusters are identified by unique gene signatures. (A) Normalized gene counts for scRNA-seq gene signatures. (B) Mouse molecular signature database hallmark gene set enrichment analysis for scRNA-seq gene signature clusters.

**Fig. S3:**
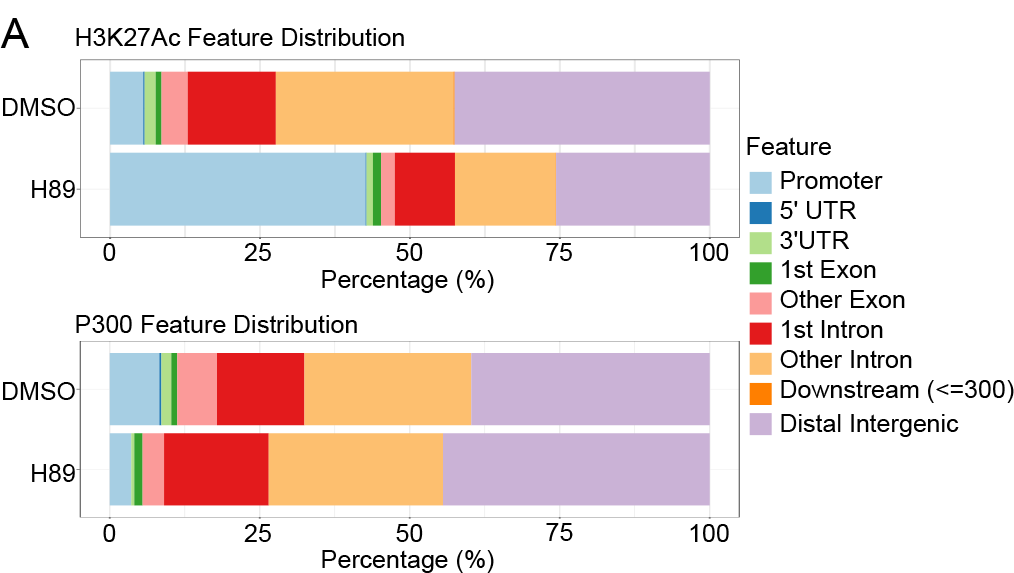
Genomic feature distribution reveals shifts in H3K27Ac and P300 binding sites following PKA inhibition. (A) top: Genomic feature distribution of H3K27Ac in DMSO or H89 treated As4.1 cells. bottom: Genomic feature distribution of P300 in DMSO or H89 treated As4.1 cells.

**Fig. S4:**
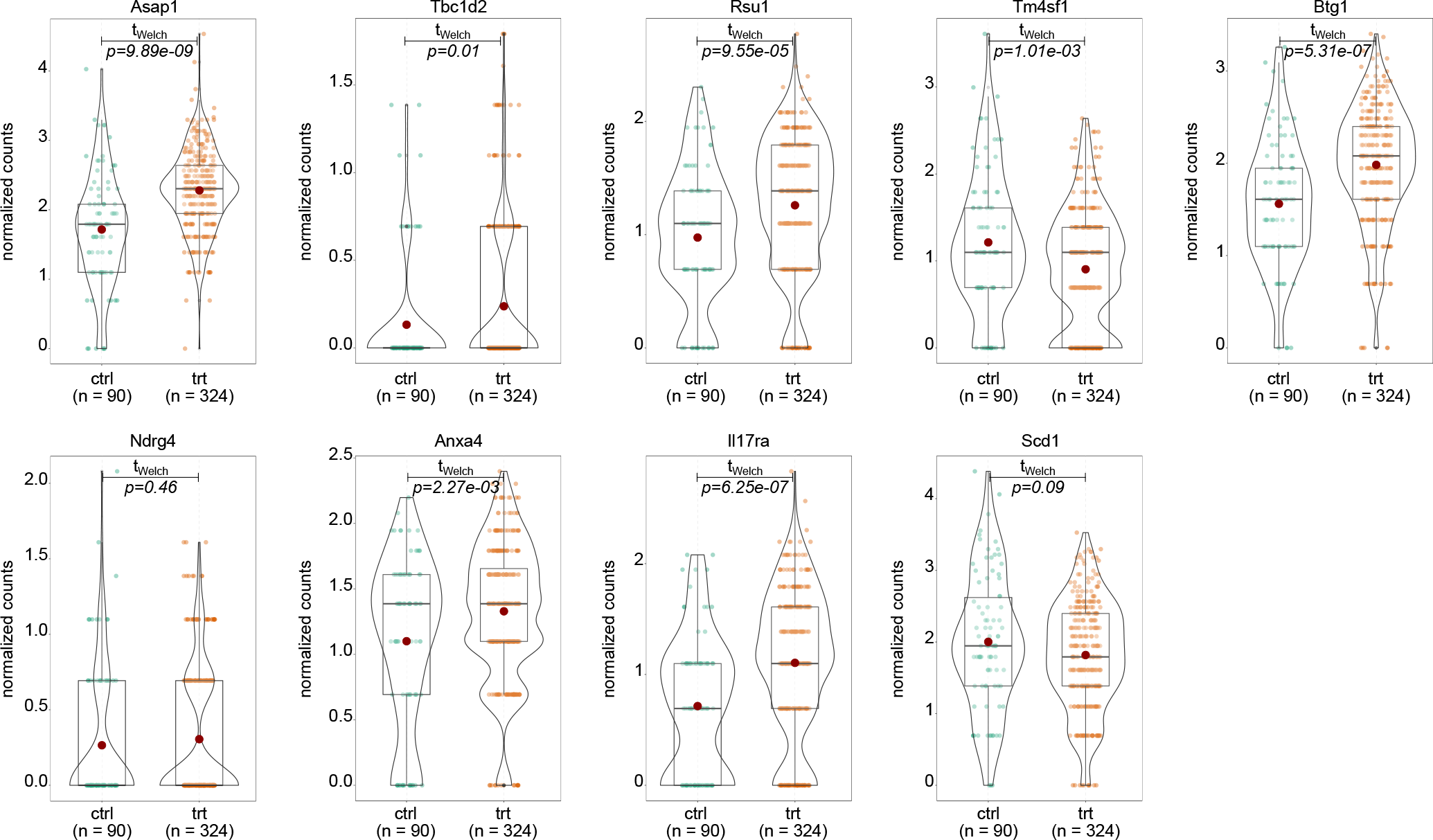
Normalized expression of restricted set of PKA inhibition centric genes within the lowest Ren1 expressing scRNA-seq cluster.

**Fig. S5:**
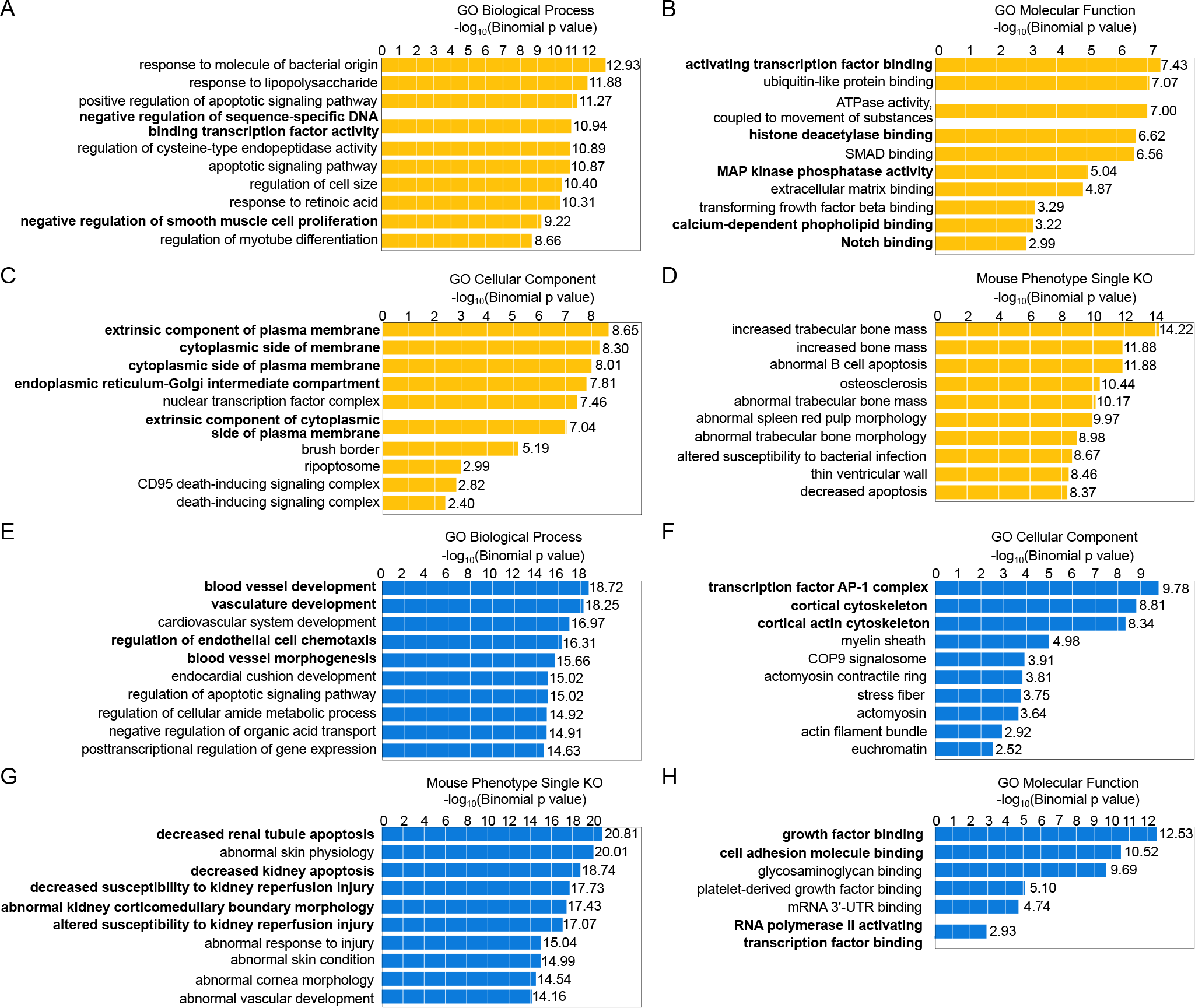
Genomic region enrichment analysis. There is a concerted loss of a contractile, smooth muscle and epithelial phenotype among the increased accessibility dynamic peaks as seen in region enrichment among (A) Biological Processes, (B) Molecular Function, (C) Cellular component, and (D) single knockout (KO) phenotypes. In relation, dynamic regions with decreased accessibility were associated with vascular development, aging, and proliferative capacity as seen across (E) Biological Processes, (F) Molecular Function, (G) Cellular component, and (H) single knockout (KO) phenotypes

